# Swimming direction of the glass catfish is responsive to magnetic stimulation

**DOI:** 10.1101/2020.08.13.250035

**Authors:** Ryan D. Hunt, Ryan C. Ashbaugh, Mark Reimers, Lalita Udpa, Gabriela Saldana De Jimenez, Michael Moore, Assaf A. Gilad, Galit Pelled

## Abstract

Several marine species have developed a magnetic perception that is essential for navigation and detection of prey and predators. One of these species is the transparent glass catfish that contains an ampullary organ dedicated to sense magnetic fields. Here we examine the behavior of the glass catfish in response to static magnetic fields which will provide valuable insight on function of this magnetic response. By utilizing state of the art animal tracking software and artificial intelligence approaches, we quantified the effects of magnetic fields on the swimming direction of glass catfish. The results demonstrate that glass catfish placed in a radial arm maze, consistently swim away from magnetic fields over 20 µT and show adaptability to changing magnetic field direction and location.

## Introduction

Throughout evolution, organisms have developed unique strategies to become more competitive in their environment. One unique adaptation is the ability to sense magnetic fields, i.e., magnetoreception. While animals like salmonids, pigeons, eels and sea turtles use magnetoreception to migrate over thousands of kilometers [1–10], non-migratory fish species have also shown evidence of magnetoreception [11, 12]. The glass catfish, is also known to be sensitive to the Earth’s magnetic field [13, 14].

The glass catfish (*Kryptopterus vitreolus)* is found in slow moving fresh water in Southeast Asia, ranges from 31.4-64.6 mm in length and is transparent except for their organ packed head [15]. The glass catfish has historically been of interest to a wide range of scientific disciplines, including circulation [16], cell line establishment [17] and electroreception [18, 19]. The interest in this species’ magnetoreception has recently been reignited due to its potential to be part of a synthetic system that will allow remote, magnetic control, of neural function [20].

New advances in molecular biology have brought new tools to identify the mechanisms by which this species respond to electromagnetic fields. Recently, we have discovered a gene (electromagnetic perceptive gene (EPG)) that is expressed in the glass catfish’s ampullary organ and is specifically activated in response to magnetic stimuli. This genetic-based magnetoreception has a great potential as a neuromodulation technology and as a valuable tool to study neural behavior from the molecular to network levels [20–22]. However, the mechanism by which magnetoreception manifests and functions is not well understood [23–29].

This work was designed to characterize the natural behavior of glass catfish in response to magnetic fields. This understanding may lead to improved engineering of magnetic-receptive modulation and sets a foundation for a new magnetically sensitive animal model. We capitalized on new concepts of artificial intelligence as well as traditional video tracking algorithms to quantify how glass catfish respond to magnetic stimulation with high spatial and temporal resolution.

## Methods

All animal procedures were conducted in accordance with the NIH Guide for the Care and Use of Laboratory Animals and approved by the Michigan State University Institutional Animal Care and Use Committee.

Glass catfish were imported from Thailand to the United States and housed in a standard 30-gallon fish tank with a 12-hour day/night cycle with water maintained at 27 degrees and provided with an enriched environment. After arriving from Thailand, fish were acclimatized for 6 weeks before starting experiments. Water quality was checked daily for ammonium, nitrite, and nitrate. Fish were fed a diet of fresh hatched brine shrimp twice per day. All experiments were conducted between the hours of 11 am and 4 pm to eliminate behavioral changes due to feeding [30] and light cycle.

The Y-maze’s arms were 60 cm long and 10 cm wide with a central area of 10×10×10 cm (AnyMaze, San Diego Instruments, CA). One week prior to starting experiments all fish were transitioned from their tanks to the radial Y-maze where they lived for the duration of each experiment (2 weeks). Enrichment materials were carried over from permanent housing to Y-maze, rearranged daily and removed prior to starting experiments. A five-gallon water change was done weekly in the Y-maze with mature water to control water quality and reduce the impact of any unknown pheromones. The same fish were used for all experiments. The magnetic stimulus and the sham stimulus were placed 10 cm from the end of an arm, inside of the maze. Each trial was recorded for 30 minutes by overhead cameras while the experimenter was out of the room. Each trial was repeated four times for each condition for a total of 24 trials. In order to negate the effects of Earth’s intrinsic magnetic field, the location of the magnet was rotated during the changing location experiment: Arm 1 was oriented in the south west direction, Arm 2 in the northern direction and Arm 3 in the south east direction.

A permanent Neodymium Rare Earth Magnet with a horizontal magnetic flux of 577 mT at the magnet’s surface was placed 10 cm from the end of one of the Y-maze arms. The strength of the magnetic field induced by the magnet was calculated by COMSOL (Figure 1). A sham stimulus was made from plastic and aluminum foil, with similar dimensions to the magnet. All recordings were analyzed by a radial-maze tracking software written in Matlab by Delcourt *et al.* [31]. Delcourt *et al.* software is available at the following address: https://github.com/sjmgarnier/projectRadial. Videos were taken originally in AnyMaze format and converted to .mp4. The Matlab program then created a background image by taking an average of 100 frames. Fish location was determined by subtracting the background image from each frame, remaining pixels with a grey scale value higher than threshold were given a value of 1, continuous pixels with a value of 1 were labeled as a fish. The spatial resolution of all videos was 3.57 ± .52 pixels/cm and were recorded at 30 frames per second. Data is reported as mean ± standard deviation. In a separate set of experiments, a single fish was selected from the school and placed in the center of the Y-maze. Two trials of magnet and sham conditions were conducted: one set of trials was used for computer training, and the other for analysis using the trained software. DeepLabCut [32, 33] was used to track the location of a single fish in the Y-maze. DeepLabCut code is available at the following address: https://github.com/DeepLabCut/DeepLabCut.git. The program was initially trained on 20 frames and ran through 1,300,000 iterations. After initial training, outlier frames were extracted and re-labeled. The program was re-trained with the addition of the outlier frames for 100,000 iterations. Retraining was done three times until visual inspection and expected error were satisfactory (±15 pixels). DeepLabCut results were extracted using an output CSV file from the code, which provides the x,y coordinates of the tracked fish for every frame of the video. This file was exported to R-Studio and plotted as a standard scatter plot and overlaid on a Y-maze diagram. Tracked videos are two minutes and fifty seconds long (5,100 frames).

**Figure 1.**
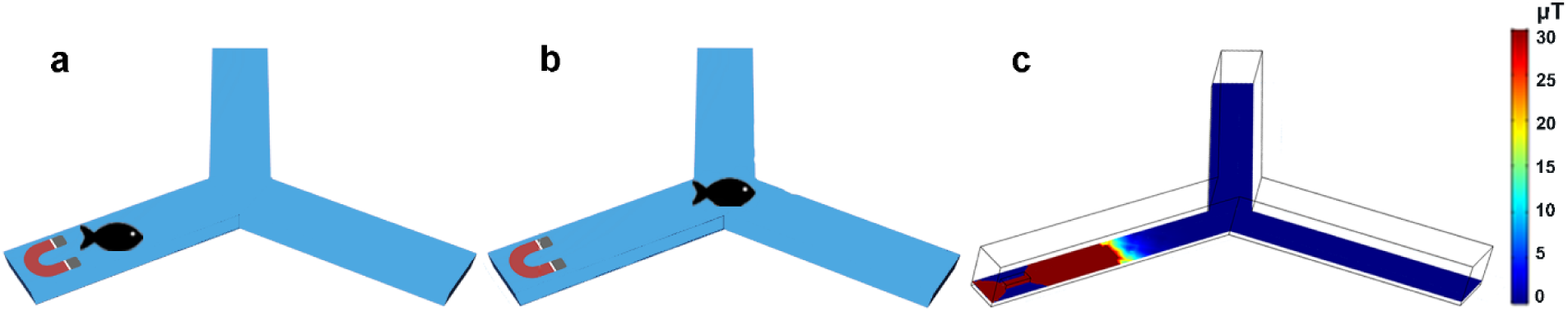
Diagram of the experimental set up. **a)** Constant location of stimulus-The magnet was always placed in the same arm and the fish were barricaded randomly in one of the three arms. **b)** Changing location of stimulus-The magnet was randomly placed in one of the three arms, and fish were always barricaded in the center of the Y-maze. **c)** COSMOL stimulation depicting the strength of the magnetic field induced by the magnet.

## Results

### Constant location of stimulus

We characterized fish behavior as a response to a magnetic stimulation that was consistently present at the same location. In these sets of experiments the magnet was always placed in Arm 1 of the Y-maze. The initial location of the fish (n=13) was changed randomly to one of the Y-maze’s three arms. Each trial was repeated four times for each arm for a total of 24 trials. The number of fish present in Arm 1 (Arm with magnet) was significantly lower than the number of fish in the other two arms in the first minute (Arm 1-Magnet, 1.24 ± .1.16; Arm 2 & 3-No Magnet, 5.16 ± 3.23; P= < .0005, unpaired T-Test), after 5 minutes (Arm 1-Magnet, 1.14 ± .91; Arm 2 & 3-No Magnet, 5.37 ± 3.38; P = < .0005, unpaired T-Test), and over the entire recording that lasted 30 minutes (Arm 1-Magnet, 1.08 ± .96; Arm 2 & 3-No Magnet, 5.02 ± 3.49; P = < .0005, unpaired T-Test). When the initial fish location was also in Arm 1, the school immediately swam away from that arm and stayed away. In contrast, when the sham stimulus was placed in Arm 1, the fish did not show any preference to any of the three arms after 1 minute, (Arm 1-Sham, 2.26 ± .1.85; Arm 2 & 3-No Sham, 3.81 ± 3.42; P= < .0005 unpaired T-Test), after 5 minutes, (Arm 1-Sham, 2.34 ± 1.95; Arm 2 & 3-No Sham, 3.96 ± .3.12; P= < .0005 unpaired T-Test), and after 30 minutes, (Arm 1-Sham, 2.07 ± 1.82; Arm 2 & 3-No Sham, 4.13 ± 3.12; P= < .005 unpaired T-Test). Between experiments we also see that fish spend significantly less time in Arm 1 when the magnet is present compared to when the sham is present at all time points (p<.0005, unpaired T-Test). These experiments showed that glass catfish prefer to avoid swimming in water with a magnetic strength over 20 µT (Figure 1c and 2).

**Figure 2.**
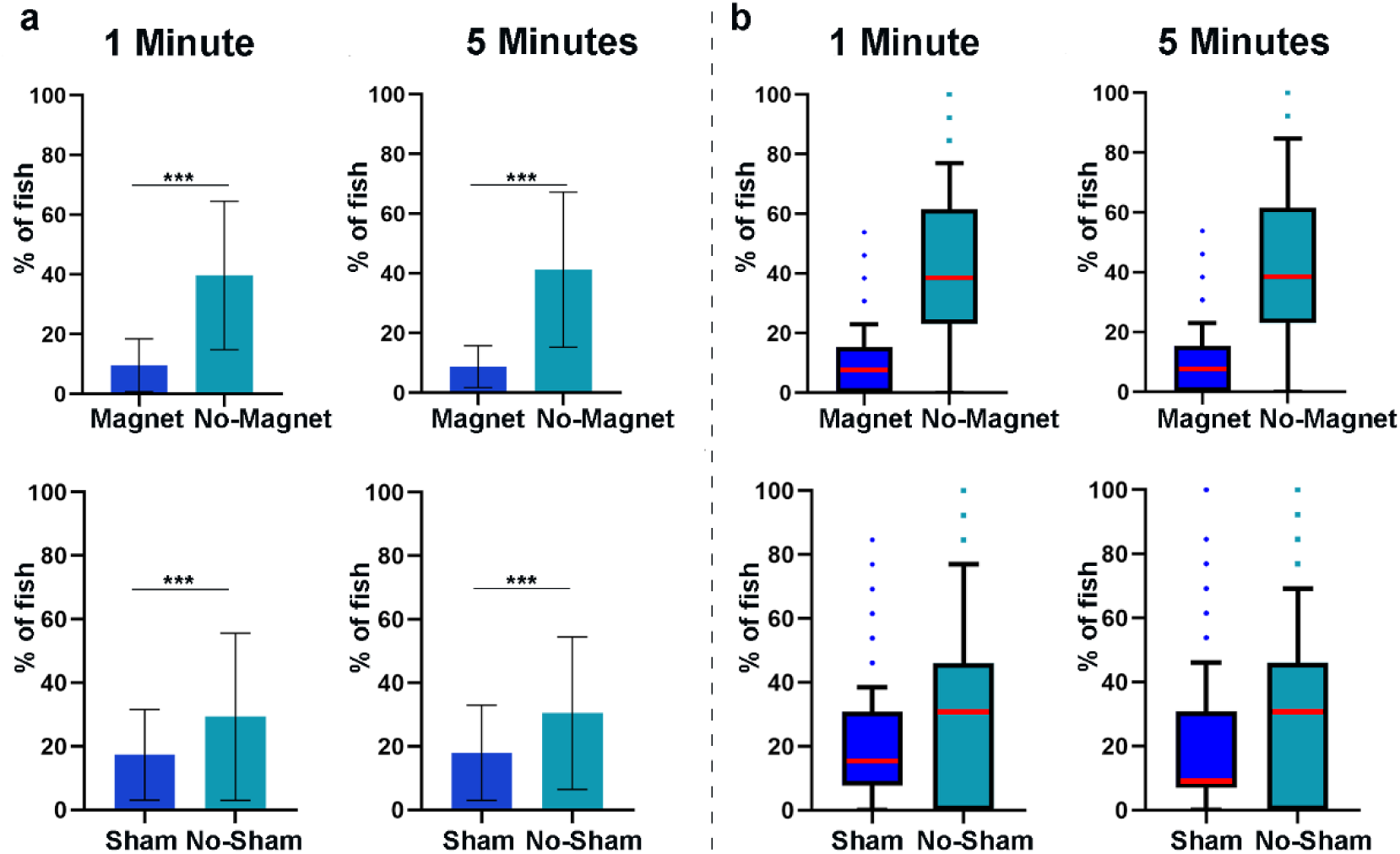
Constant location of stimulus results. **a)** The average number of fish in each arm over the first minute and first five minutes of experiment. **b)** Distribution of fish over first minute and five minutes of experiment, red line indicates median value, outliers shown are >95% CI. The magnet and sham object were kept in Arm 1 across all trials. Results indicate that regardless of the initial location of the fish, they tend to avoid Arm 1 when magnet was present compared to sham (*** p<.0005, unpaired T-Test).

### Changing location of stimulus

We then sought to determine if fish behavior changed with the location of the magnetic stimulation. In this set of experiments the fish school (n=12) was barricaded in the middle of the Y-maze, and the magnetic or sham stimuli were placed randomly in one of the arms (Figure 1b). After barricade removal the fish swam away from the magnet and explored the two other maze-arms. In line with previous experiments, fish spent significantly less time in an arm when the magnet was present compared to when the sham stimulus was present (Arm 1-Magnet, 1.58 ± .95, Arm 1-Sham, 3.76 ± 1.97; Arm 2-Magnet, 1.39 ± .99, Arm 2-Sham, 3.37 ± 1.29; Arm 3-Magnet, 6.63 ± .1.29, Arm 3-Sham, 8.53 ± 1.11; P= < .0005, Unpaired T-Test). Figure 3 shows the number of fish present in each arm within the first 5 minutes of recording.

**Figure 3.**
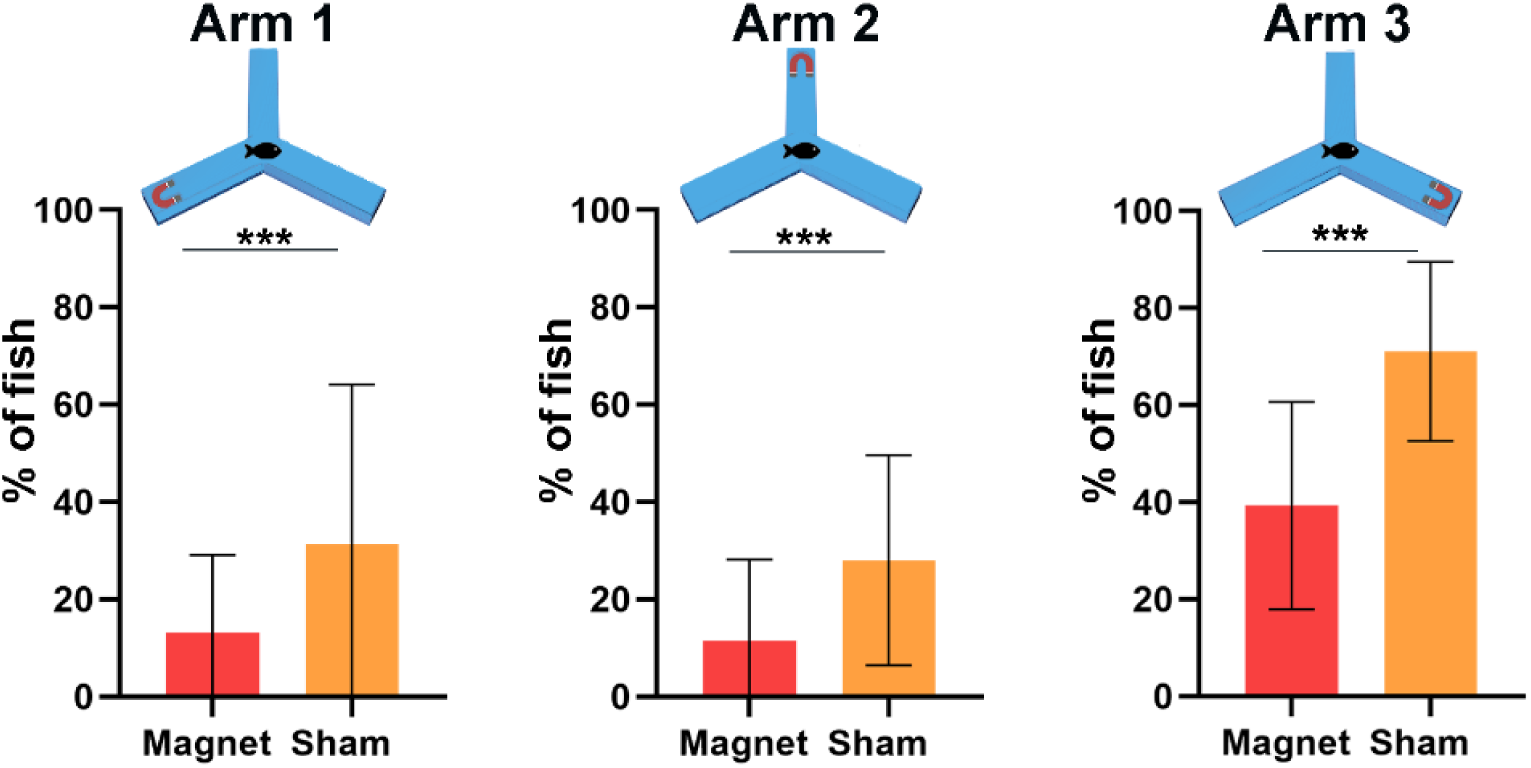
The average number of fish in each arm over the five minutes in changing location of stimulus experiments. There is a significant decrease in number of fish between magnetic stimulus and sham in every arm of the y-maze (*** p<.0005, unpaired T-Test).

For the purpose of individual swim pattern analysis, one fish was placed in the middle of the Y-maze. Once the barricade had been removed the fish swam across two arms where no magnet was present but exhibited a clear avoidance from the arm containing the magnet. We used a state-of-the-art artificial intelligence (AI) approach, DeepLabCut[32, 33] to track the fish’s swimming path. DeepLabCut was successfully trained on a single fish with an error of less than 4.2 cm or 15 pixels. Figure 4 shows that magnetic stimulation results in an individual fish swimming away from the magnet immediately after barricade removal. Consistent with previous results, sham stimulus did not induce an avoidance behavior.

**Figure 4.**
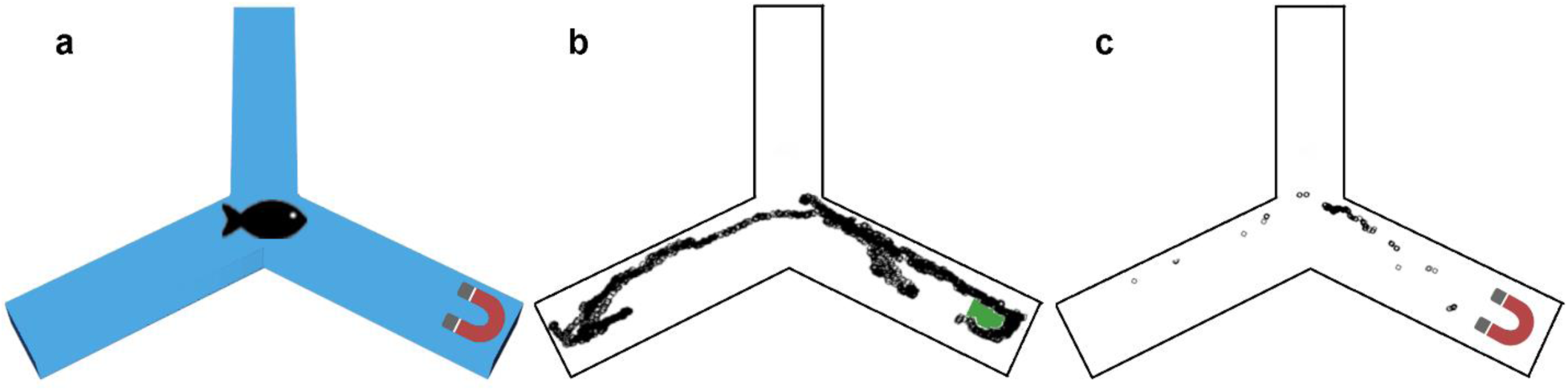
DeepLabCut tracking of a single fish. **a)** Experimental set up. **b)** Sham, **c)** magnet. The fish position is shown here over every frame for 170 s (5,100 frames). Indivudual values were exported and plotted using R and overlaid on a Y-maze diagram.

## Discussion

Several marine species have developed a magnetic perception that is useful in navigation and the detection of prey and predators [6, 34], for review see [35–37]; animals such as sharks and platypus, use magnetoreception for prey detection [38–42]. Others, like ants [43], use this sense for predator avoidance and the nematode, *Caenorhabditis elegance* uses magnetoreception for vertical navigation in soil [44]. Even cattle have been shown to align themselves with electromagnetic pulses [45]. The glass catfish, is a transparent fish found in slow moving rivers of Southeast Asia where visibility is low[15, 17]. Until recently, the transparent glass catfish was commonly identified as *Kryptopterus bicirrhis* but is now known to be *Kryptopterus vitreolus* [15]. It appears plausible that in these types of conditions, magnetic perception is an advantageous trait to conserve. However, the mechanisms allowing magnetic sensation remain largely unknown [26, 37]. We have used state-of-the-art software based on artificial intelligence object tracking algorithms to characterize glass catfish behavioral response to magnetic fields. The results indicate that glass catfish consistently swim away from magnetic fields over 20 µT and show adaptability to changing magnetic field direction and location. In addition, our results show that this magnetic avoidance behavior is not influenced by school behavior. We have previously demonstrated that the modulatory effects of magnetic stimulation on mammalian cells transfected with EPG was induced by magnetic fields of 50 mT [21]. However, in this experiment we see that fish are significantly more sensitive to magnetic fields than transfected cells, with fish starting to exhibit avoidance behavior at ∼20 µT (Figures 2 and 3). While the pathway by which EPG modulates calcium channels is unknown [20, 21] it is possible that there are accessory proteins unknown to the authors, which amplify magneto-sensitivity in glass catfish. Currently, we can only evoke a cellular response by using strong magnetic fields in culture (> 50mT). However, the Earth’s magnetic field is only 30 µT-60 µT, and yet, is readily detected by glass catfish. One of the major challenges in EPG’s development as a neuromodulatory technology is the attenuation of magnetic fields over distance. If the biological amplification properties of the glass catfish is uncovered, this technology could be used to treat deep brain afflictions without the need for surgery.

Using AI such as DeepLabCut can be transformative to animal behavior studies. Using this method, we could follow the swimming pattern of an individual fish, over thousands of frames with extremely high spatial and temporal resolution. Another advantage is the machine learning components of AI. The more trials run through DeepLabCut, the more efficient and accurate it becomes at tracing animals in similar situations. However, the transparency of glass catfish caused detection difficulties with DeepLabCut and recording hardware when rapid movement caused insufficient contrast between the fish and Y-maze. In Figure 4b, during the sham stimulus the fish swam at a gradual pace throughout the maze. However, during magnetic stimulus the fish tend to stay in one area then dash to the end of an arm and back. During these rapids movements there was not enough contrast for the software to detect the fish. Once the fish slowed down and the contrast was restored, the tracking became accurate.

We have established that the glass catfish has unique magnetic field sensing capabilities that position it as a valuable model to study magnetoreception in animal species. The cellular mechanisms allowing this capability remains to be determined. We have already identified and cloned the EPG from glass catfish. But is this the only magnetic-sensitive protein? Does it work with other proteins to amplify and modulate its activity? Do other animal species that have been shown to be sensitive to magnetic fields have similar proteins? This animal model can provide unprecedent preparation to address these questions. By characterizing the behavior of glass catfish, we are now working towards developing a fish with a knock-out in the EPG gene. This will elucidate if there are additional genes associated with magnetic responses and will facilitate the development of the next generation of additional magnetic sensing molecular tools.

## Acknowledgements

This work was supported by National Institutes of Health grants R01NS072171 and R01NS098231.

## Authors Contributions

RDH and GP designed all experiments and wrote the manuscript. RDH and GSDJ performed experiments. RDH, RCA, MM, MR, AG and GP analysed the data. RCA and LU performed the magnetic measurements, and related analysis.

